# Kinesin-1 motility traced by an activity-based precipitating dye

**DOI:** 10.1101/2020.08.12.247544

**Authors:** Simona Angerani, Eric Lindberg, Nikolai Klena, Christopher K. E. Bleck, Charlotte Aumeier, Nicolas Winssinger

## Abstract

Kinesin-1 is a processive motor protein that uses ATP-derived energy to transport a variety of intracellular cargoes toward the cell periphery. As tracks for cargo delivery, kinesin-1 uses a subset of microtubules within the dense microtubule network. It is still debated what defines the specific binding of kinesin-1 to a subset of microtubules. Therefore, the ability to visualize and monitor kinesin transport in live cells is critical to study the myriad of functions associated with cargo trafficking. Herein we report the discovery of a fluorogenic small molecule substrate for kinesin-1 that yields a precipitating dye. The activity of kinesin-1 thus leaves a fluorescent trail along its walking path and can be visualized without loss of signal due to diffusion. Kinesin-1 specific transport of cargo from the Golgi appears as trails of fluorescence over time.

## Introduction

Microtubules (MTs) are polymers of α and β tubulin that are involved in several functions in cells. Although the majority of MTs emanates from the centrosome,^1^ the main non-centrosomal MT organizing center is represented by the Golgi apparatus.^2^ The minus-end of the MT is anchored at the MT organizing center and the dynamic plus-end orientated towards the cell periphery.

Motor proteins, such as kinesins and dyneins, are ATPases that bind to MTs and walk along them in response to cargo binding.^3^ Kinesin-1 is a member of the kinesin family that transports cargoes to the cell periphery walking on MTs towards their plus-end.^4^ Among others, kinesin-1 is interacting with both pre-Golgi and Golgi membranes and it is involved in Golgi-to-ER and ER-to-Golgi trafficking.^5,6^ Kinesin-1 is autoinhibited and only functionally active once bound to a cargo during Golgi-to-ER transport. The motion of kinesin-1 occurs preferentially on a subset of modified, long lived MTs, such as acetylated and detyrosinated MTs.^7,8,9^ The transport activity of kinesin-1 can be inhibited by Taxol, a drug which stabilizes and changes the MT structure.^10^

To date, techniques to track motor proteins in cells have relied on antibodies, quantum dots or on engineered versions of the motors bearing fluorescent tags;^11-16^ these techniques require sample treatment (fixation and staining) or manipulation (transfection). Moreover, these techniques stain total protein content, irrespectively of their motility. Only about 30 % of kinesin-1 is active in cells;^17^ this makes it extremely difficult to study kinesin-1-GFP movement along MTs within the strong background of immotile kinesin-1-GFP in transfected cells.^7^

QPD is a quinazolinone-based precipitating dye developed to easily visualize enzymatic activity *in cellulo*.^18,19^ Accordingly, QPD has been used to design fluorogenic reporters of phosphatase (PO_4_^-^ derivative),^20^ protease (ester derivative)^21^ and H_2_O_2_ (boronic acid derivate)^22^ or catalysis (azide^23^ or picolinium^24^ derivative), and these substrates have been used to label a number of organelles and cytoskeletal elements.^20, 23^ QPD fluorescence derives from an excited-state intramolecular proton transfer (ESIPT)^25^ between the phenolic group and the quinazolinone. Functionalization of the phenol frees the aryl moiety out of planarity with the quinazolinone, which dramatically reduces its aggregation and precipitation; derivatization with a polar group renders these molecules water soluble.

Herein, we report the discovery of a QPD derivative (QPD-OTf) that acts as an activity-based fluorogenic substrate for kinesin-1 by producing a precipitating fluorescent dye along its walking path on MTs. The phenolic moiety is functionalized with a triflate group that renders the molecule soluble in aqueous buffer and non-fluorescent. Biochemical experiments showed that kinesin-1 and MTs were sufficient to yield fluorescent crystals and that inhibition of kinesin1’s ATPase activity reduced the formation of fluorescent crystals. Docking studies support the binding of the activity-based substrate in the nucleotide binding site, aligning the triflate leaving group with the gamma-phosphate group of ATP. In live cells, the crystals are centered in the Golgi apparatus and radially elongate towards the cell periphery, following the path tracked by kinesin-1 motion on MTs. To our knowledge, this is the first time that the native transport activity of kinesin-1 is visualized *in cellulo* without external modifications.

## Results

### QPD-OTf forms crystals in living cells

Taking advantage from the large applicability of QPD-based profluorophores, we envisioned the synthesis of a QPD derivative, QPD-OTf, (Fig. 1) initially designed to be responsive to superoxide, for the visualization of oxidative stress *in cellulo*. In analogy with a reported fluorescein derivative,^26^ the trifluoromethanesulfonate ester should be activated enough to undergo nucleophilic attack by O_2_^.-^ affording the free phenol. Surprisingly, QPD-OTf was found not to be responsive to O_2_^.-^ *in vitro*, with no precipitation observed.

**Fig. 1.**
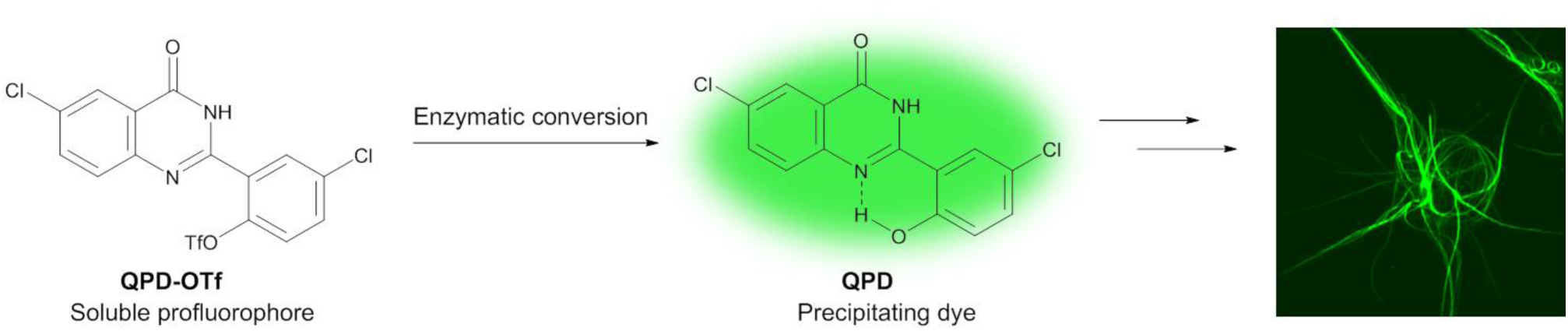
QPD precipitation and crystal formation in cells.

QPD-OTf treatment of zymosan stimulated RAW264.7 cells caused dotted fluorescent precipitate after 10 min that evolved into complex filamentous crystals within 1 hour (Supplementary Fig. 1). The QPD crystal is an extended, aster-like fluorescent crystals expanding throughout the entire cell and even able to deform the cell membrane (Fig. 1, Supplementary Fig. 2). While overnight exposure of 10-20 μM QPD-OTf induces cell death, temporary exposure of up to 4 hours followed by fresh media replacement, preserves cell viability almost completely (Supplementary Fig. 3) and even dissolved the crystal over time. Since the fluorescent signal can only arise from a QPD displaying an uncaged phenol, we hypothesize that the triflate caging moiety must be removed inside the cell upon enzymatic activity. The observation of these crystals across multiple cell lines from different mammalian species (RAW264.7, HeLa, MCF-7, HEK293, U2OS, PTK2) shows that this activity is conserved and not restricted to a specialized cell line (Supplementary Fig. 2).

**Fig. 2.**
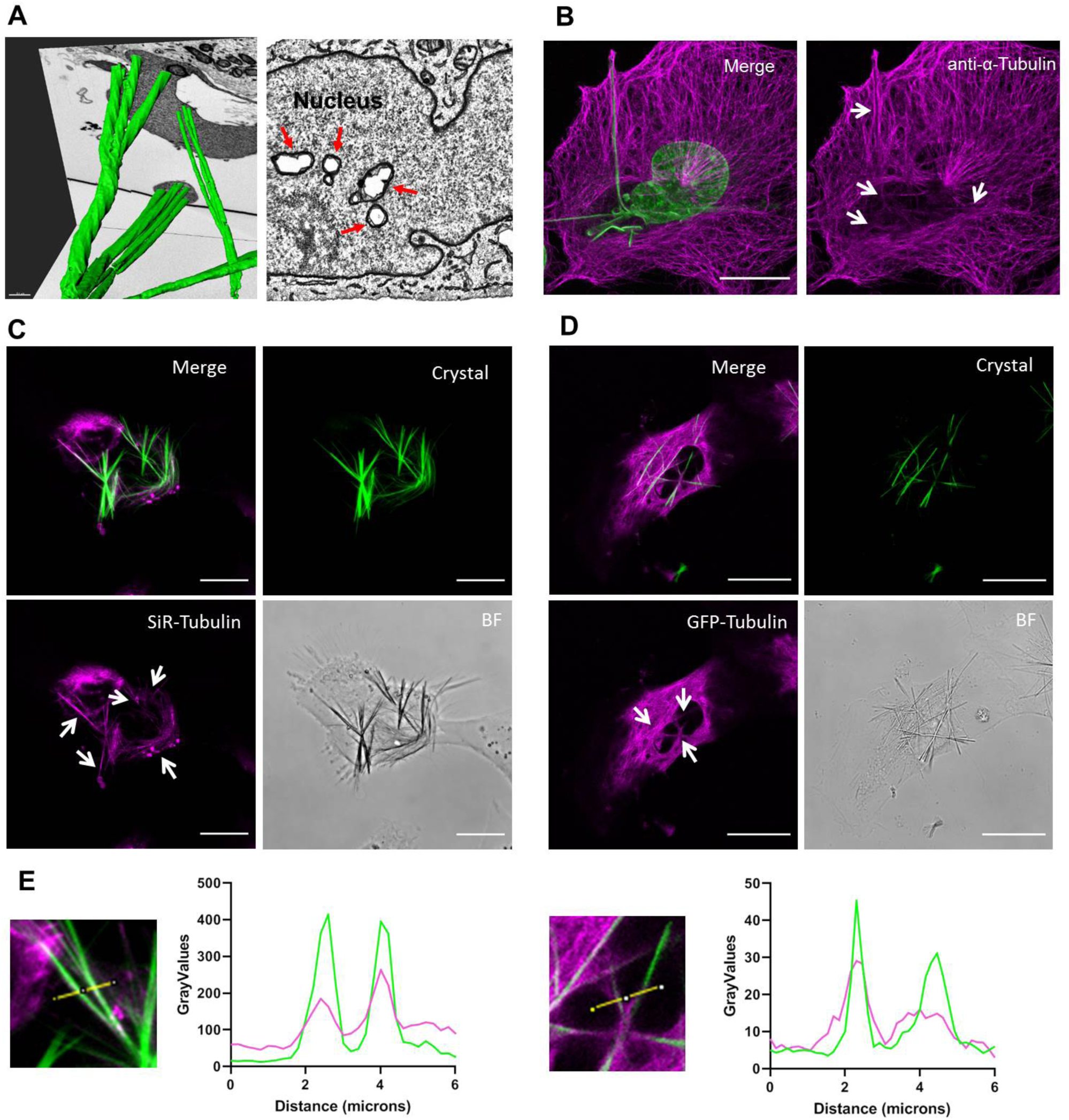
Crystal analysis in relation to cellular elements. A) FIB-SEM 3D-reconstruction of crystal inside HeLa cell (left); FIB-SEM cross section of HeLa cell containing crystals spanning through the nucleus (right); red arrows indicate the crystal sections. B) Tubulin immunostaining in fixed U2OS treated with QPD-OTf; white arrows indicate colocalization with crystals (anti α-tubulin: magenta; crystal and DAPI: green). C) Tubulin staining in live U2OS cells treated with QPD-OTf; white arrows indicate colocalization with crystals (SiR-Tubulin: magenta; crystal: green). D) Live cell imaging of PTK2-GFP-Tubulin treated with QPD-OTf; white arrows indicate colocalization with crystals (GFP-tubulin: magenta; crystal: green). E) Plot profiles of tubulin channel and QPD channel for images shown in C and D. Scale bar 20 μM.

**Fig. 3.**
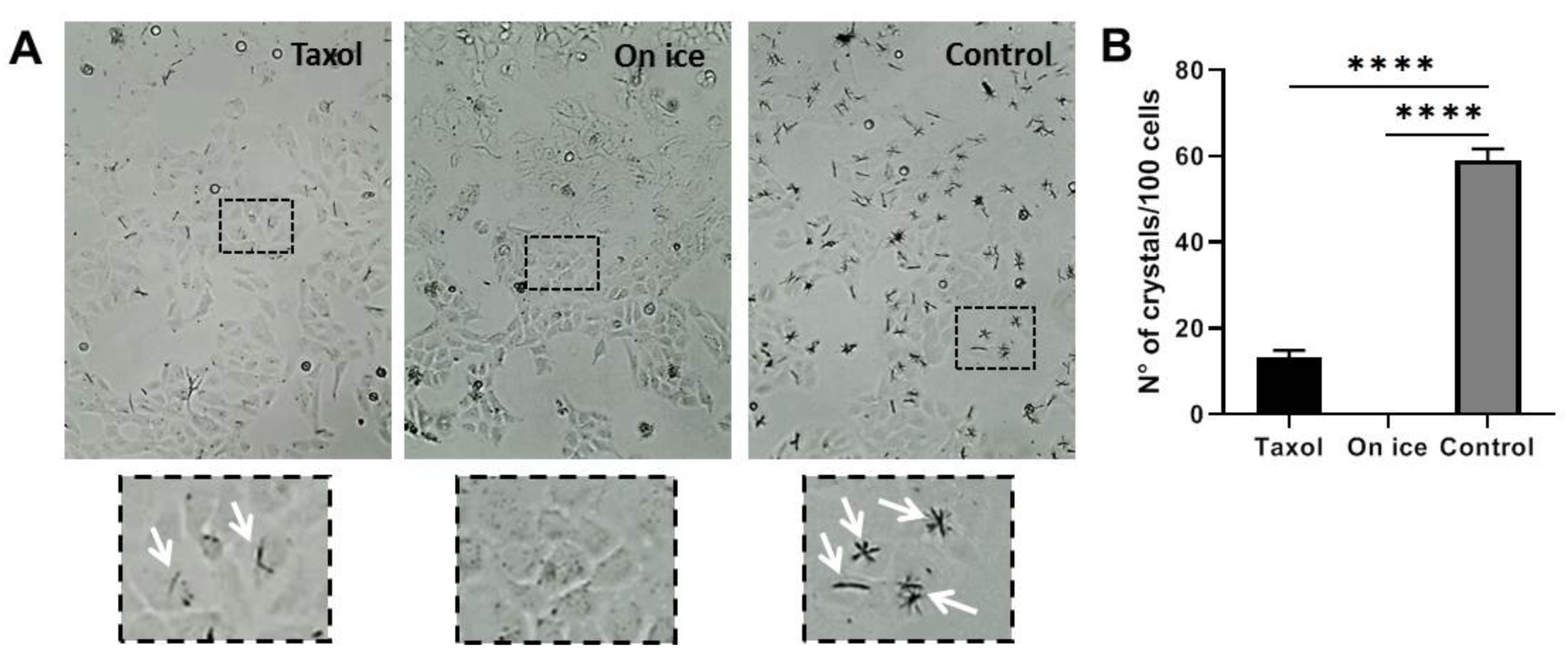
Effect of microtubules stabilization/depolymerization on crystal formation in U2OS live cells. A) Taxol 1 μM 1 hour + QPD-OTf 20 μM 4 hours at 37 °C (left); on ice 1 hour + QPD-OTf 20 μM 4 hours on ice (center); QPD-OTf 20 μM 4 hours at 37 °C (right), and zoom of cells defined in the black square (bottom); white arrows indicate crystals. B) Quantification of number of crystals for conditions reported in A.

FIB-SEM analysis of HeLa cells treated with QPD-OTf (20 μM, 4 hours) showed that the crystals have a well-defined organization, with rotational symmetry order 3-like structure (Fig. 2A, left), and hexagonal cross-section, whose size varies from 100 to 700 nm (Supplementary Fig. 4). The rigidity of the crystals is sufficient to induce membrane deformation as crystals spanning through the nucleus were observed (Fig. 2A, right). We also noted that the crystal has a clear nucleation center (Supplementary Fig. 4) which spurred us to further investigate the triggering mechanism behind the crystal formation.

**Fig. 4.**
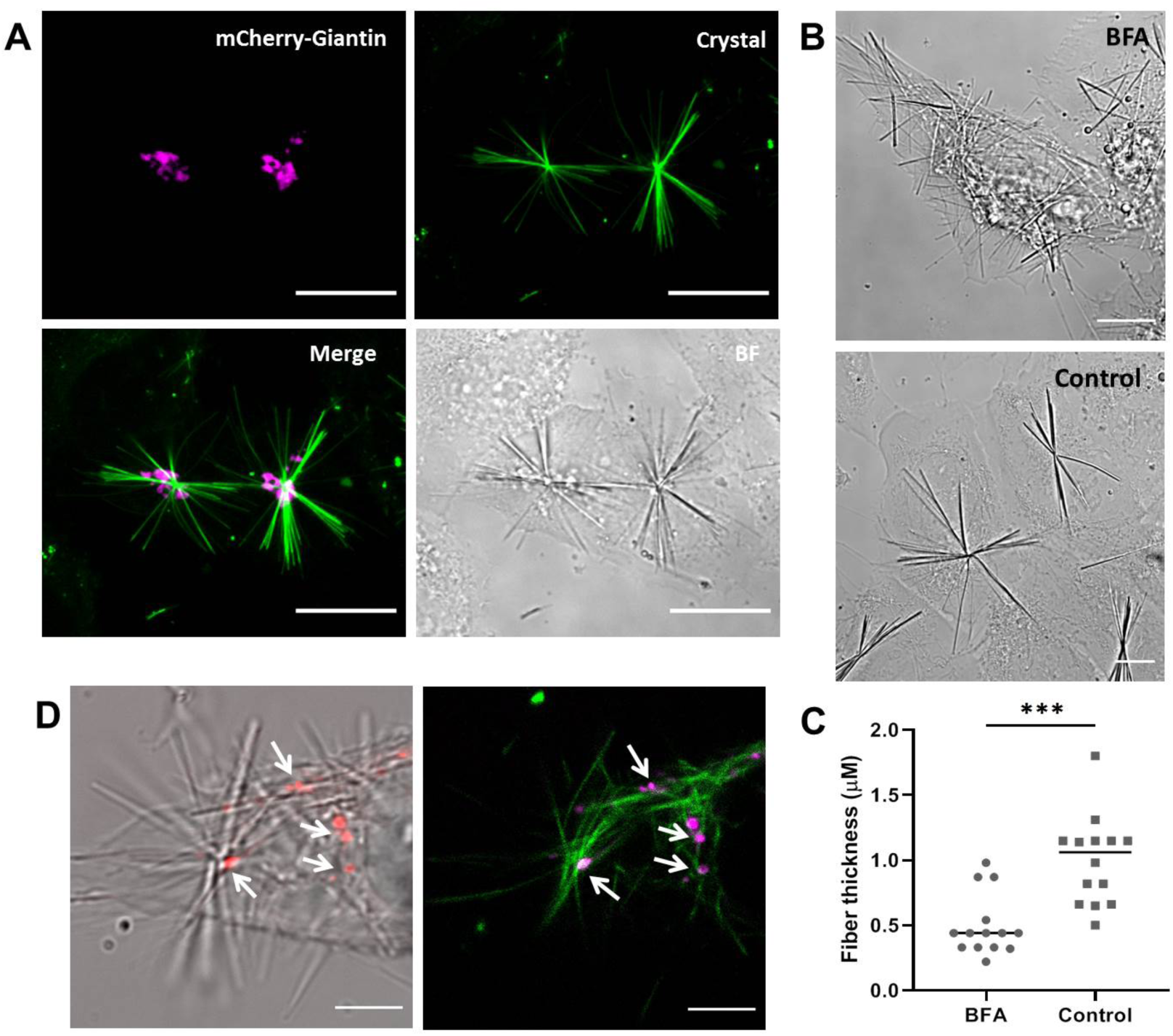
Correlation between the center of crystals and Golgi elements. A) U2OS-mCherry-Golgi + QPD-OTf (20 μM 3 hours). Crystals (green), mCherry-Giantin (magenta). Scale bar 20 μM. B) BFA effect on crystal morphology and location. BFA treated cells (20 μM BFA 4 hours + 20 μM QPD-OTf 2.5 hours (top). Control (20 μM QPD-OTf 2.5 hours) (bottom); scale bar 10 μM. C) Quantification of images reported in B. D) Localization of crystals and Golgi vesicles after BFA treatment; Golgi (red) crystals (bright field) (left); Golgi (magenta) crystals (green) (right); arrows indicate centers of crystal; scale bar 5 μM.

### Crystals co-localize with MTs

Many crystals are localized at the cell center, spanning with their filamentous nature throughout the cell. Due to their organization and architecture we thought that the enzymatic activity generating QPD-crystals might be linked to the actin or MT cytoskeleton. Labeling the cytoskeleton after QPD-OTf treatment showed clear colocalization between the crystal and the MT network and only marginal correlation with actin (Fig. 2, Supplementary Fig. 5). In fixed cells, immunostaining of α-tubulin showed alignment of crystal fibers along MT bundles (Fig. 2B). We observed that only a distinct subset of the MT network seemed to colocalize with the crystal. Live cell imaging by expressing GFP-tubulin in Ptk2 cells, or staining MTs with SiR-Tubulin,^27^ a Taxol based fluorogenic dye, confirmed the colocalization of the crystal with a subset of the MT network (Fig. 2C,D,E).

**Fig. 5.**
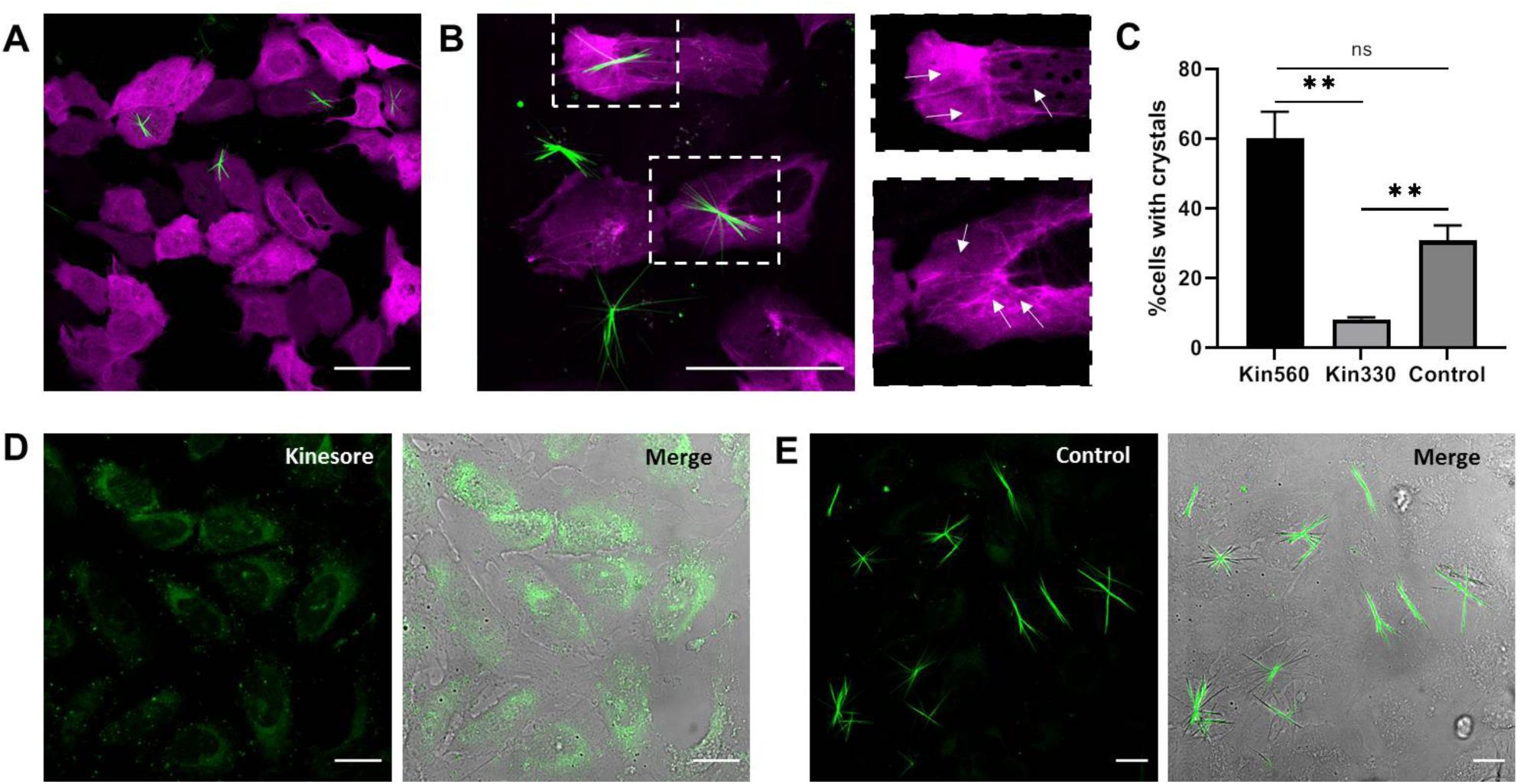
The role of kinesin-1 in crystal frormation. A) PTK2-GFP-Tubulin transfected with kin330 plasmid and treated with QPD-OTf (20 μM, 2.5 hours); green: crystals, magenta: kinesin. B) PTK2-GFP-Tubulin transfected with kin560 plasmid and treated with QPD-OTf (20 μM, 2.5 hours) (left); zoom of highlighted boxes, arrows indicate stabilized MTs correlating with crystals (right); green: crystals, magenta: kinesin. Scale bar: 50 μm. C) Quantification of crystal formation in transfected cells *vs* control. D) U2OS treated with kinesore (100 μM) in Ringer’s buffer + QPD-OTf (20 μM); green: QPD fluorescence. E) Control conditions for experiment reported in D (QPD-OTf 20 μM, 2 hours in Ringer’s buffer); green: crystals. Scale bar 20 μM.

Not all MTs within the cellular network have the same dynamical properties, and we wondered if MT dynamics was linked to crystal formation upon QPD-OTf treatment. In order to test our hypothesis, we altered MT dynamics and studied the impact on crystal formation. To stabilize MTs we treated U2OS cells with 1μM Taxol, followed by addition of 20 μM QPD-OTf. Even after incubation for 4 hours, only very few crystals were observed compared to the control (Fig. 3A). Taxol treatment reduced the crystal formation by 75% (Fig. 3B). Moreover, the few crystals we observed in the Taxol-treated sample were much thinner than in the control (Fig. 3 zoom). Then, we completely depolymerized the MT network by cold treatment, followed by 20 μM QPD-OTf incubation for 4 hours. In this case, no crystals were observed (Fig. 3). The almost complete absence of crystals with both treatments suggests that the integrity and physiological dynamic of MTs are substantial requirements for crystal development.

### QPD crystals track MTs originating from the Golgi apparatus

To identify the origin of the specific localization of the crystal within a subset of the MT network, we focused on the nucleation site of the crystals. In fact, most of the cells in interphase show a single crystal, originating close to the nucleus. This raises the possibility that the nucleation site of the crystal overlaps with the nucleation site of MTs. MTs nucleate mainly from MT organizing centers located close to the cell center. However, the most prominent MT organizing center, the centrosome, did not co-localize with the triggering site for QPD precipitation (Supplementary Fig. 6). Therefore, we investigated another MT organizing center: the Golgi apparatus. The Golgi is known to be involved in MT nucleation,^2,28^ and to be a key player in the secretory pathway.^29^

**Fig. 6.**
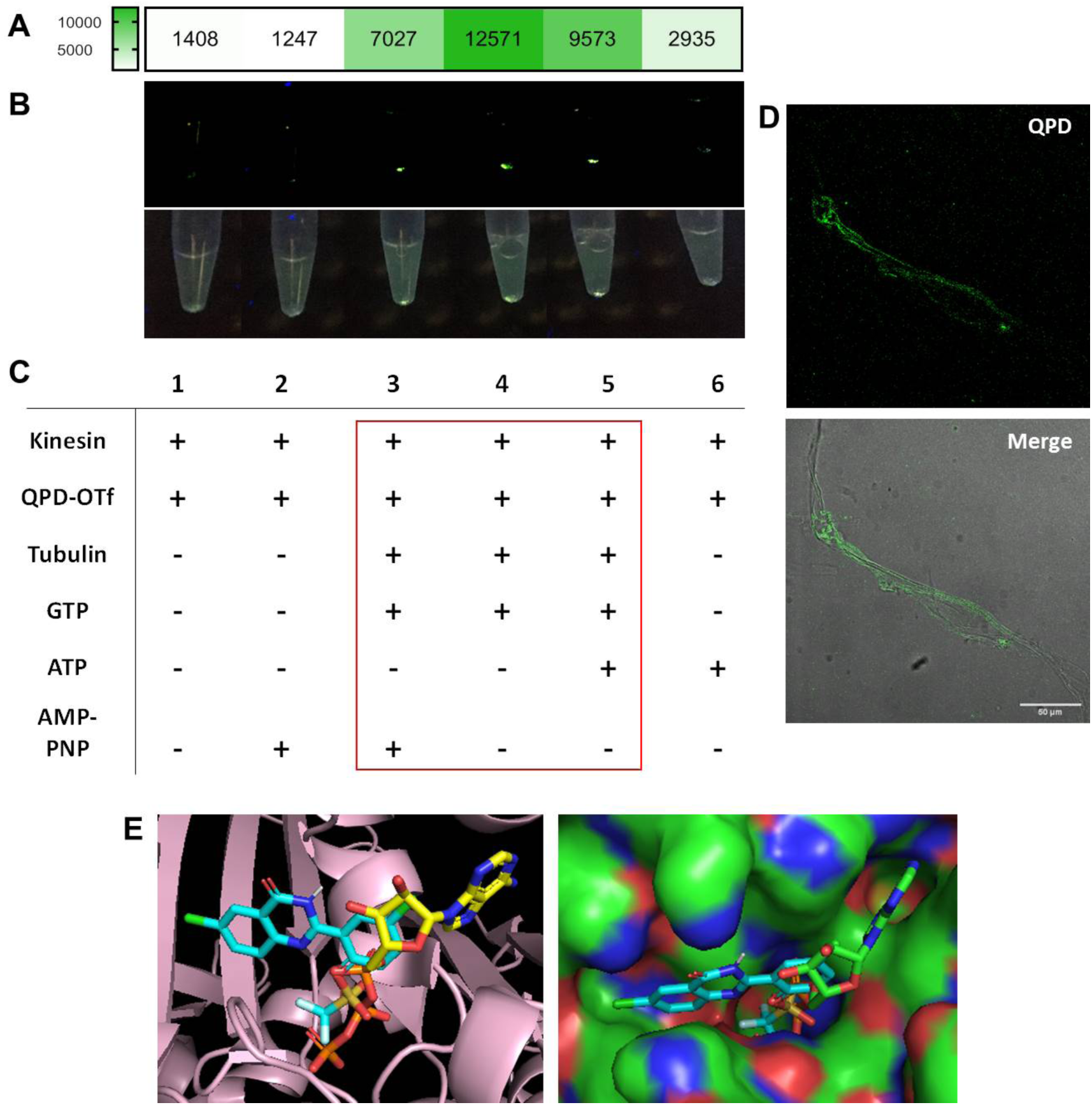
*In vitro* and computational studies of QPD-OTf. A) In vitro precipitation of QPD; intensity map of emitted light by samples 1-6 under 366 nm excitation; intensity values are expressed as grey values from the green channel of an RGB picture acquired with a smartphone camera. B) Samples under 366 nm light: green channel (top); original picture (bottom). C) Table of conditions for the samples reported in B; red square indicates the samples that gave detectable QPD-fluorescence. D) Confocal imaging of fluorescent filaments contained in sample 4. Scale bar 50 μm. E) Docking of QPD-OTf into ATP binding site of kinesin-1. Left: QPD-OTf (cyan), ATP (yellow), kinesin-1 (pink); right: QPD-OTf (cyan), ATP (green), kinesin-1 (polarized surface).

To assess if the MT organizing center at the Golgi triggers QPD precipitation, we visualized the Golgi in U2OS cells by transfecting them with mCherry-Giantin. Confocal fluorescence microscopy revealed that the crystals are nucleated at the Golgi apparatus (Fig. 4A). The observation was confirmed by transfecting mCherry-Giantin in a stable expressing PTK2-GFP-tubulin cell line with sequential treatment of QPD-OTf (Supplementary Fig. 7). The MT organizing center of the Golgi, together with its transport activity could therefore play a key role in determining the selective transformation of QPD-OTf to QPD in a specific cellular location.

In order to further investigate the role of the Golgi apparatus in the formation of QPD crystals, we studied the effect of Brefeldin A (BFA), an inhibitor of Golgi trafficking. BFA impairs the function of Golgi, resulting in its fragmentation.^30,31^ While the Golgi apparatus appears as a compact complex in U2OS interphase cells, treatment with 20 μM BFA showed the expected scattered Golgi fragments (Supplementary Fig. 8). QPD-OTf addition to BFA treated cells resulted in thinner crystals (fiber thickness reduced by 58%) (Fig. 4B, C) with multiple foci of origin instead of only one as in control cells (Fig. 4B). Although the Golgi was fragmented, the nucleation site of the crystal remained co-localizing with the Golgi (Fig. 4D). This shows that the Golgi apparatus is intimately linked to crystal formation and that modifications of the Golgi structure correlate with crystal morphology and location.

### Purified MTs are not sufficient to generate crystals *in vitro*

Having established that MT dynamics is necessary for the development of crystals, and that the Golgi apparatus, known as a MT organizing center, dictates the location of the crystals, we assessed whether pure MT polymerization is sufficient to generate a crystal *in vitro*. To this end, we tried to precipitate QPD on Taxol stabilized MTs, or on dynamic MTs elongating from stabilized seeds. No crystal formation could be observed and no fluorescence of QPD was detected in our *in vitro* TIRF assay, even after 2 hours. Thus, we reasoned that the conversion of QPD-OTf to QPD crystals must be triggered by an enzymatic event that is closely related to and dependent on the MT network, but an activity that is not essential for MT elongation. With these considerations, we directed our attention to motor proteins since these proteins move cargoes along MTs in an energy dependent manner.

### QPD-OTf conversion to QPD depends on kinesin-1 motility

The plus end directed motor protein kinesin-1 transports cargo from the Golgi to the ER and its enzymatic activity might be responsible for the conversion of QPD-OTf to QPD with ensuing crystal formation. We therefore genetically modified kinesin-1 activity in cells and analyzed the effect of kinesin-1 activity on crystal formation. Cells were treated with QPD-OTf after transfection with kin330-GFP or kin560-GFP, two truncated versions of kinesin-1 fused to GFP displaying no ability to walk on MTs, and constitutively active walking on MTs respectively.^32,33^ In cells transfected with the immotile kin330 the number of crystals was reduced by 87% compared to non-transfected cells (Fig. 5A, Supplementary Fig. 9, and Fig. 5C). The residual formation of some crystals could be attributed to the activity of endogenous wildtype kinesin-1. Although, over-activity of kinesin-1 in cells transfected with kin560-GFP did not further increase crystal formation (Fig. 5B, Supplementary Fig. 9). It was possible to correlate the crystal filaments to the signal of kin560-GFP on MTs (Fig. 5B zoom). It is also noteworthy that despite the concentration of kin560-GFP and broad distribution, fluorescent crystals are only seen on specific tubulin axis.

To further investigate the effect of kinesin-1 activity on crystal formation, we tested the effect of adding kinesore, a small molecule kinesin-1 activator.^34^ In cells, kinesin-1 is inactive and only gets activated upon cargo binding.^35, 36^ Kinesore interacts with kinesin-1 at the kinesin light chain-cargo interface (K_i_ = 49 μM for aiKLC2^TPR^: SKIP^WD^ complex), mimicking the effect of cargo binding and resulting in kinesin-1 activation. The enhanced motion causes profound rearrangement of the MT network. We found that addition of 100 μM kinesore to U2OS cells, followed by incubation with 20 μM QPD-OTf inhibited the formation of crystals, yet generated some diffuse QPD fluorescence (Fig. 5 D-E). This diffused fluorescence as a result of kinesore treatment is attributed to the overactivity of kinesin-1, with a motor activity that is no longer coupled to its endogenous localization or regulation. This result strengthens the involvement of kinesin-1 activity in QPD formation and corroborates the results observed with kin330 transfection. Moreover, the fact that QPD formation is observed in the presence of kinesore suggest that kinesore does not compete directly with QPD-OTf binding.

### Kinesin-1 forms QPD crystals *in vitro*

In a cell, multiple proteins can interact and show enzymatic activity. To pin down if kinesin-1 is the candidate to convert QPD-OTf to QPD crystals we analyzed a reconstituted *in vitro* system with purified proteins. We tested the crystal formation under several conditions in presence of kinesin-1, +/- tubulin, MT, ATP, GTP, AMP-PNP in BRB buffer. Samples containing both kinesin-1 and MTs had a strong QPD fluorescence (Fig. 6A-C, samples 3,4,5), with the most intense signal deriving from the sample containing QPD-OTf, kinesin, tubulin and GTP (sample 4). In addition, filamentous structures were observed in the MT/kinesin/QPD-OTf samples. Confocal microscopy confirmed that the filamentous-QPD structures were fluorescent (Fig. 6D). In presence of a non-hydrolysable analogue of ATP (AMP-PNP), where kinesin-1 is motility is reduced while bound to MTs,^37, 38^ we observed lower levels of fluorescent precipitate (Fig. 8A-C, sample 3). This reduced signal is consistent with our *in cellulo* observation where kinesin-1 motor activity is required for QPD-OTf conversion. The presence of ATP also slightly reduced the formation of the precipitate (Fig. 8A-C, sample 5). Collectively, this shows that kinesin-1 converts QPD-OTf to QPD and suggests that QPD-OTf binding is competitive (directly or allosterically) with ATP. These results suggest a potential interaction between QPD-OTf and the ATP binding site of kinesin-1; the ATP-ase activity of the kinesin-1 motor domain might be serving as enzymatic activity responsible for the triflate cleavage. The fact that fluorescent crystals are observed along the filaments in the absence of ATP suggests that QPD-OTf can act as a substrate for kinesin-1.

### QPD-OTf as a substrate analogue of ATP

Taken together, the cellular and biochemical data show the dependence of crystal formation upon kinesin-1 motion on MTs. Since kinesin-1 exploits ATP hydrolysis to propel its motor domain processively on MTs,^39^ and that ATP is not required for crystal formation while AMP-PNP reduces crystal formation, we hypothesized that QPD-OTf acts as a substrate analog. We performed molecular docking of QPD-OTf into the ATP binding pocket of the kinesin kinesin-1 motor domain. We calculated the fitting into human kinesin-1 in the ATP state (PDB: 3J8Y) using Autodock Vina.^40^ The best pose offered a calculated binding energy of −8.3 kcal/mol. Superposition of this binding pose with ATP showed that the triflate overlaps with the hydrolyzed phosphate of ATP (Fig. 6E). Based on the structural similarities between QPD and ispinesib^41^, an allosteric Eg5 inhibitor which also has a chloroquinozolinone moiety, we also performed docking calculations for QPD-OTf in the nucleotide binding site of both Eg5 and kin-1. QPD-OTf shows good pose correlation with Ispinesib (Supplementary Fig. 10) and good affinity for Eg-5 in its nucleotide binding pocket (−8.2 kcal/mol); however, this pose positioned the triflate towards the solvent, making the triflate hydrolysis impossible (Supplementary Fig. 11). Docking studies with kinesin-1 indicated less favorable binding (−5.8 kcal/mol) in the allosteric site (Supplementary Fig. 12). Collectively, these docking studies support a direct hydrolysis of the triflate of QPD-OTf and provide a rational for the selectivity of kinesin-1 over Eg5 and kin-1.

## Discussion

Small molecule fluorophore conjugates have been a powerful approach to track a protein of interest and the development of fluorogenic probes for live-cell imaging of the cytoskeleton, for example, have empowered cellular biology studies.^27^ Alternatively, fluorogenic probes have been designed to report on a given enzymatic activity by introducing a masked fluorophore as a leaving group in an enzymatic reaction, thus acting as activity-based fluorescent reporter.^42^ While this approach has been extremely productive to image hydrolytic enzymes, such as protease and glycosidase, with a specific substrate recognition and broad tolerance for the leaving group, there are no examples reported for motor proteins. The discovery of a fluorogenic substrate (QPD-OTf) to image kinesin-1 in live cells shows that it is possible. Moreover, the hydrolysis of a phenolic triflate represents a new modality for activity-based probes. This substrate is particularly attractive for a motor protein since its fluorescent product precipitates and leaves a bright fluorescent trail along the path traveled by kinesin-1.

Until now it was difficult to trace native kinesin-1 activity in cells. Kinesin-1-GFP expression at native level results in a high fluorescent background of inactive kinesin-1-GFP and it is therefore impossible to distinguished which microtubules are used for transport.^15^ Complex experimental setups have been developed, like tracing microtubule dynamics *in vivo*, fixing cells and adding purified tagged kinesins to map which microtubules are likely to be used for transport. Our dye for the first time shows a possibility to trace native kinesin-1 activity live in a cell without any modification or fixation. The development of QPD-OTf opens the possibility to map the usage of a subset of microtubules within the dense and dynamic microtubule network.

In summary, we report an activity-based substrate for kinesin-1 yielding a bright precipitate in response to kinesin-1 activity along MTs. Based on the kinesin-1’s transport activity from the Golgi, fibers are observed as a function of time, developing from foci at the Golgi. The center of the crystals reflects the location of Golgi elements; the number of crystals per cell and their thickness correlates with Golgi compactness/fragmentation. The crystal formation is sensitive to kinesin-1 motility; kinesin-1 inhibitors disrupt the formation of the crystals. In addition, the presence of MTs is required to generate QPD fluorescence *in vitro*. The biochemical data and docking studies support an ATP competitive mechanism involving QPD-OTf binding to the nucleotide pocket and acting as a substrate resulting in triflate hydrolysis. The resulting QPD product precipitates to form a bright fluorescent fiber along the microtubules used by kinesin-1. QPD-OTf staining is compatible with live cell imaging; the possibility to dissolve the crystals in cell media after staining provides a non-destructive method to visualize the motion of kinesin-1 on Golgi derived MTs.

## Methods

### Synthesis of QPD-OTf

4-chloro-2-formylphenyl trifluoromethanesulfonate (see supplementary materials for synthetic procedure of this compound) (300 mg, 1.04 mg, 1 eq), 2-amino-5-chlorobenzamide (177 mg, 1.04 mmol, 1 eq), *p*-toluensulfonic acid (59 mg, 0.3 mmol, 0.3 eq) were suspended in EtOH (2.6 mL). The mixture was heated to reflux and stirred for 2 hours. The the mixure was cooled to 0 °C and 2,3-Dichloro-5,6-dicyano-1,4-benzoquinone (236 mg, 1.04 mmol, 1 eq) was added portion wise; the mixture turns to green, then to brown with a white precipitate. The suspension was diluted with cold EtOH (5 mL) and centrifuged. The white solid was washed twice with cold EtOH and dried under vacuum. The crude residue was purified by flash chromatography on silica gel (Pentane/EtOAc 9:1 to 6:4). Fractions containing the desired product were collected and dried under reduced pressure to afford QPD-OTf (360 mg, 80% yield). Exact mass: 437.946. LC-MS (ESI^+^): RT= 2.92 min. m/z found: 438.94 [M+H]^+. 1^H NMR (400 MHz, Acetone-*d*6): δ 11.69 (s, 1H), 8.18 (dd, *J* = 2.5, 0.5 Hz, 1H), 8.13 (d, *J* = 2.6 Hz, 1H), 7.89 (dd, *J* = 8.8, 2.5 Hz, 1H), 7.85 (dd, *J* = 8.9, 2.6 Hz, 1H), 7.82 (dd, *J* = 8.8, 0.5 Hz, 1H), 7.68 (d, *J* = 8.9 Hz, 1H). ^13^C NMR (101 MHz, Acetone-*d*6): δ 161.12, 149.43, 148.04, 146.44, 135.74, 134.91, 133.57, 132.29, 130.97, 130.23, 126.10, 125.55, 123.89, 120.99, 117.81. ^19^F NMR (282 MHz, Acetone-*d*6): δ −75.10. See Supplementary Fig. 13 for LC-MS traces and Supplementary Fig. 14, 15, 16 for ^1^H, ^13^C and ^19^F NMR spectra.

### Crystal formation in cells

QPD-OTf (20 μM) was added to cells in DMEM (−) without additives and incubated at 37 °C, 5% CO_2_ from 2 to 4 hours. Crystals can be easily detected by a 20X objective.

### Live cell imaging of QPD-OTf treated cells

U2OS cells (2×10^5^) were seeded into 3.5 cm glass bottom dishes with 10 mm microwell (Mattek); cells were incubated in McCoy’s 5A medium at 37 °C under 5% CO_2_ in a humidified incubator for 16 hours. Then media was removed, cells were washed twice with DMEM (−) (no additives) and QPD-OTf (20 μM) was added to cells in DMEM (−) (no additives). Cells were incubated at 37 °C under 5% CO_2_ for 4 hours. After washing twice with DMEM (−) (no additives), SiR-Tubulin (1 μM) was added and cells were incubated for 45 min at 37 °C under 5% CO_2_. Cells were washed twice with DMEM (−) and imaged with a LEICA SP8 microscope.

PTK2-GFP-Tubulin cells (2×10^5^) were seeded into 3.5 cm glass bottom dishes with 10 mm microwell (Mattek); cells were incubated in alpha-MEM medium at 37 °C under 5% CO_2_ in a humidified incubator for 16 hours. Then media was removed, cells were washed twice with DMEM (−) (no additives) and QPD-OTf (20 μM) was added to cells in DMEM (−) (no additives). Cells were incubated at 37 °C under 5% CO_2_ for 2 hours. Cells were washed twice with DMEM (−) and imaged with a LEICA SP8 microscope.

### MTs stabilization with Taxol in live U2OS cells

U2OS cells (2×10^5^) were seeded into 3.5 cm glass bottom dishes with 10 mm microwell (Mattek); cells were incubated in McCoy’s 5A medium at 37 °C under 5% CO_2_ in a humidified incubator for 24 hours. Then media was removed, cells were washed twice with DMEM (−) (no additives), Taxol (1 μM) was added to cells in DMEM (−) (no additives) and cells were incubated at 37 °C under 5% CO_2_ for 1 hour. QPD-OTf (20 μM) was then added and cells were incubated at 37 °C under 5% CO_2_ for 4 hours. Cells were imaged with a 20X objective on an EVOS XL Core.

### MTs depolymerisation on ice in live U2OS cells

U2OS cells (2×10^5^) were seeded into 3.5 cm glass bottom dishes with 10 mm microwell (Mattek); cells were incubated in McCoy’s 5A medium at 37 °C under 5% CO_2_ in a humidified incubator for 24 hours. Then media was removed, cells were washed twice with DMEM (−) (no additives) and put on ice for 1 hour. QPD-OTf (20 μM) was then added and cells were incubated on ice for 4 hours. Cells were imaged with a 20X objective on an EVOS XL Core. Control cells were washed with with DMEM (−) (no additives) and incubated with QPD-OTf (20 μM) at 37 °C under 5% CO_2_ for 4 hours.

### Transient transfection with mCherry-Giantin plasmid

U2OS or PTK2-GFP-Tubulin cells (1.5×10^5^) were seeded into 3.5 cm glass bottom dishes with 10 mm microwell (Mattek); cells were incubated in culture medium at 37 °C under 5% CO_2_ in a humidified incubator for 24 hours. pSF-mCherry-SNAP-Giantin plasmid (kind gift of Riezman’s lab; University of Geneva, Switzerland) was transfected with FugeneHD reagent in Optimem (100 μL); cells were incubated at 37 °C under 5% CO_2_ for 24 hours. Cells were washed twice with DMEM (−) (no additives) and QPD-OTf (20 μM) was added to cells in DMEM (−) (no additives). Cells were incubated at 37 °C under 5% CO_2_ for 3 hours. Cells were washed twice with DMEM (−) and imaged with a LEICA SP8 microscope.

### Brefeldin A treatment in live U2OS cells

U2OS cells (2×10^5^) were seeded into 3.5 cm glass bottom dishes with 10 mm microwell (Mattek); cells were incubated in McCoy’s 5A medium at 37 °C under 5% CO_2_ in a humidified incubator overnight. Then media was removed, cells were washed twice with DMEM (−) (no additives), Brefeldin A (20 μM) was added to cells in DMEM (−) (no additives) and cells were incubated at 37 °C under 5% CO_2_ for 4 hours. Then media was replaced with fresh one containing Brefeldin A (20 μM) and QPD-OTf (20 μM) and cells were incubated at 37 °C under 5% CO_2_ for 2.5 hours. Cells were washed twice with DMEM (−) and imaged with a LEICA SP8 microscope. The same protocol was used for cells transfected with mCherry-Giantin plasmid.

### Kinesore + QPD-OTf treatment in live cells

U2OS or PTK2-GFP-Tubulin cells (1.5×10^5^) were seeded into 3.5 cm glass bottom dishes with 10 mm microwell (Mattek); cells were incubated in culture medium at 37 °C under 5% CO_2_ in a humidified incubator for 24 hours. Then media was removed, cells were washed twice with DMEM (−) (no additives), Kinesore (100 μM) + QPD-OTf (20 μM) were then added to cells in Ringer’s buffer and cells were incubated at 37 °C under 0% CO_2_ for (1.5 hours for PTK2; 2 hours for U2OS). Cells were imaged with a LEICA SP8.

### Transient transfection with kin330-GFP/kin560-GFP

U2OS cells (1.5×10^5^) were seeded into 3.5 cm glass bottom dishes with 10 mm microwell (MatTek); cells were incubated in culture medium at 37 °C under 5% CO_2_ in a humidified incubator for 24 hours. Kin330-GFP or kin560-GFP plasmid was transfected with FugeneHD reagent in Opti-Mem (100 μL); cells were incubated at 37 °C under 5% CO_2_ for 24 hours. Cells were washed twice with DMEM (−) (no additives) and QPD-OTf (20 μM) was added to cells in DMEM (−) (no additives). Cells were incubated at 37 °C under 5% CO_2_ for 3 hours. Cells were washed twice with DMEM (−) and imaged with a LEICA SP8 microscope.

### FIB-SEM

HeLa cells were grown on glass coverslips (MatTek) and incubated with QPD-OTf (20 μM) for 4 hours. HeLa cells were processed for FIB-SEM using an rOTO protocol as previously described with some modifications^43^. Samples were processed for thin resin embedding. The samples were imaged inside a Zeiss Crossbeam 540 FIB-SEM microscope and the run was setup and controlled by Atlas software (Fibics Incorporated, Ottawa, Ontario, Canada). See Supplementary material for more details.

### *In vitro* precipitation of QPD

20 μM QPD was precipitated in Eppendorf-tubes at room temperature for 6 hours in BRB80 in presence of different combinations of unlabelled 14 μM tubulin, 150 nM kinesin-1, 2.7 mM AMP-PNP, 2.7 mM ATP, 1 mM GTP. Samples were visualized under a 366 nm lamp. The content of samples containing fluorescent precipitate was imaged by a LEICA SP8 microscope.

### Molecular docking

Docking calculations were performed with Autodock Vina. Receptor (PDB structure: 3J8Y for kinesin-1, 4AP0 for Eg5) and ligand preparation were performed in AutodockTools. Results were displayed with PyMol.

## Acknowledgement

We thank Howard Riezman’s group for providing the mCherry-Giantin plasmid. We thank Paul Guichard’s group for providing useful reagents and for constructive criticism on the manuscript. The authors thank the NCCR Chemical Biology and the University of Geneva (Département de l’instruction publique) for financial support.

## Author contributions

E.L. discovered QPD-OTf and performed seminal experiments *in cellulo*. S.A. performed or contributed to all experiments. E.L. and C.B. performed FIB-SEM experiments. N.K. performed immunostaining experiments. C.A. performed TIRF microscopy and *in vitro* experiments. S.A., E.L., N.K., C.A. and N.W. designed experiments. N.W. and C.A. supervised the project. S.A., C.A. and N.W. wrote the manuscript. All the authors discussed and commented on the manuscript.

## Additional information

Correspondence should be addressed to C.A. and N.W.

