## Supplemental information for "Kinesin-1 motility traced by an activity-based precipitating dye"

Supplementary Figures........................................................S2

Supplementary Methods........................................................S9

Supplementary References........................................................S14

### Supplementary Figures

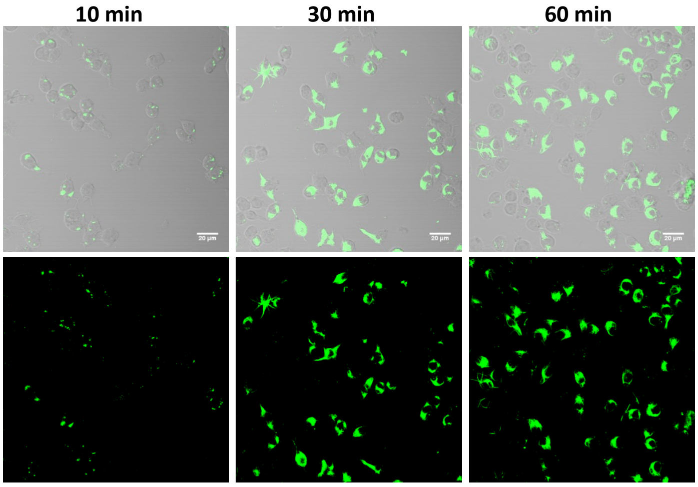


**Supplementary Figure 1. QPD-OTf in RAW246.7 cells**. QPD fluorescence (bottom) at different time points and merge images (top). Scale bar 20 µM.

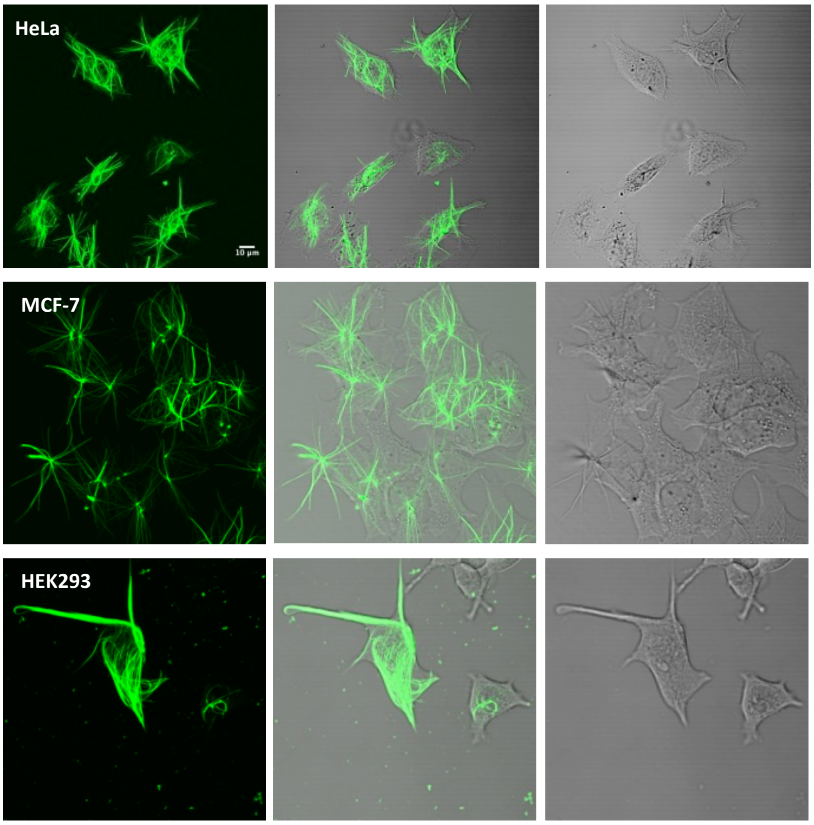


**Supplementary Figure 2**. **QPD-OTf in a panel of human cell lines.** QPD fluorescence (left); merge (centre); bright field (right).

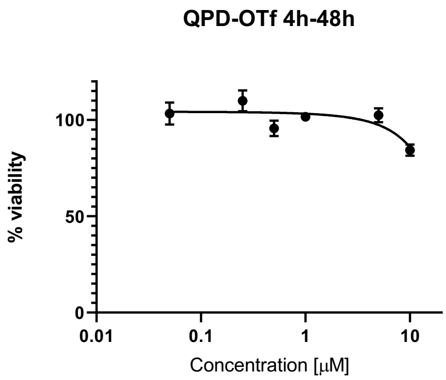


**Supplementary Figure 3. WST-1 viability assay in MCF-7 cells**. Cells were treated with QPD-OTf for 4 hours; washed, incubated for 48 hours, then WST-1 assay was performed.

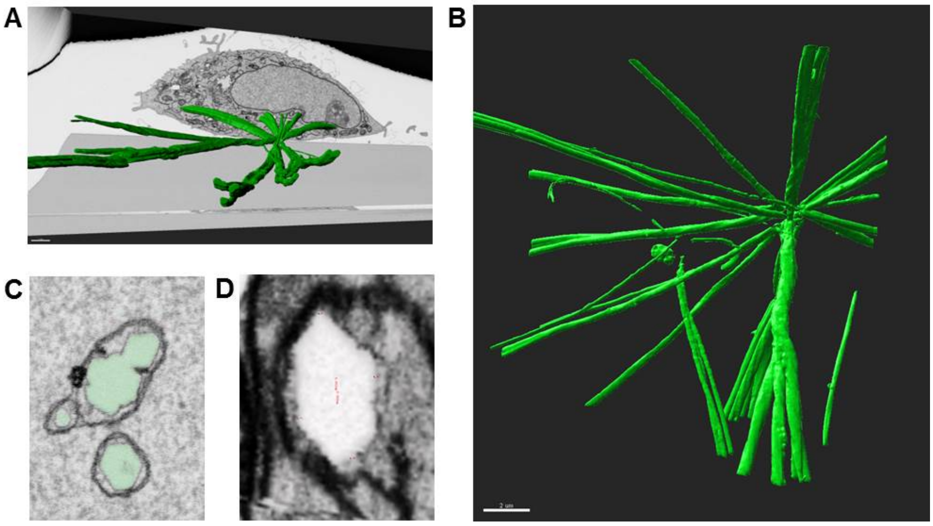


**Supplementary Figure 4. 3D reconstruction of crystals in HeLa cells from FIB-SEM analysis**. A) Crystal (green) over cell section. B) Side view of crystal. Scale bar 2 μM. C), D) Cross sections of crystal fibers.

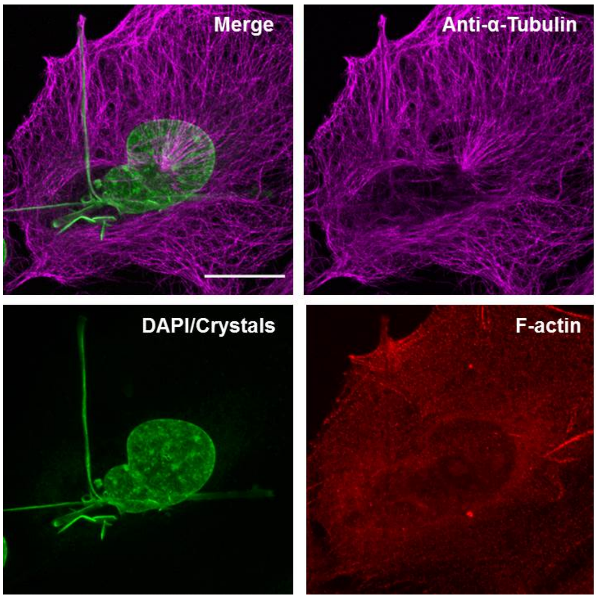


**Supplementary Figure 5. Immunofluorescence of U2OS cells treated with QPD-OTf.** Fixed U2OS treated with QPD-OTf and stained with DAPI, α-tubulin immunostaining and actin immunostaining. Green: DAPI/Crystals; magenta: anti-α-tubulin; red: F-actin.

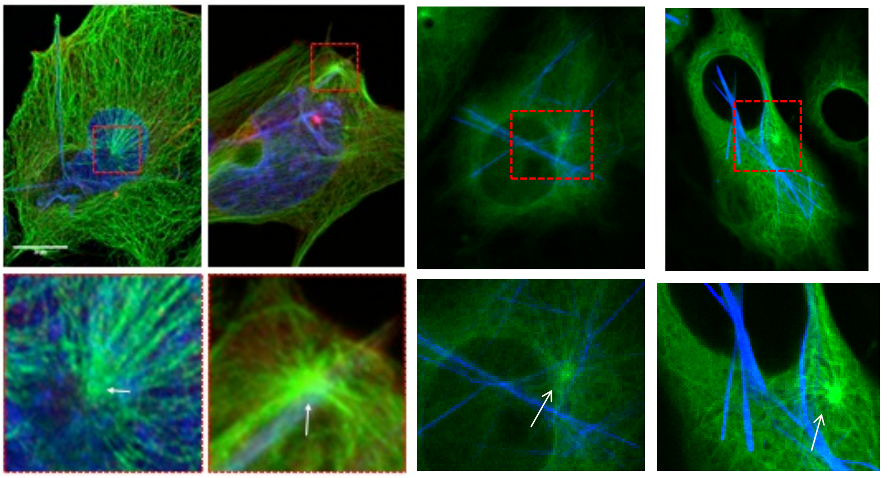

**Supplementary Figure 6. Centrosome position in QPD-OTf cells**. Green: tubulin; blue: crystal. Arrows indicate the position of the centrosome.

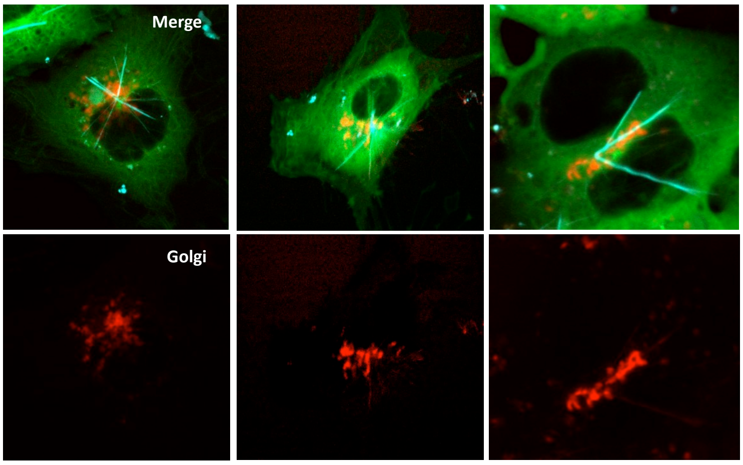


**Supplementary Figure 7. Colocalization of the center of crystals with Golgi apparatus**. PTK2-GFP-Tubulin cells were transfected with mCherry-Giantin plasmid and treated with QPD-OTf (20 µM, 2 hours). Green: tubulin; cyan: crystals; red: Golgi.

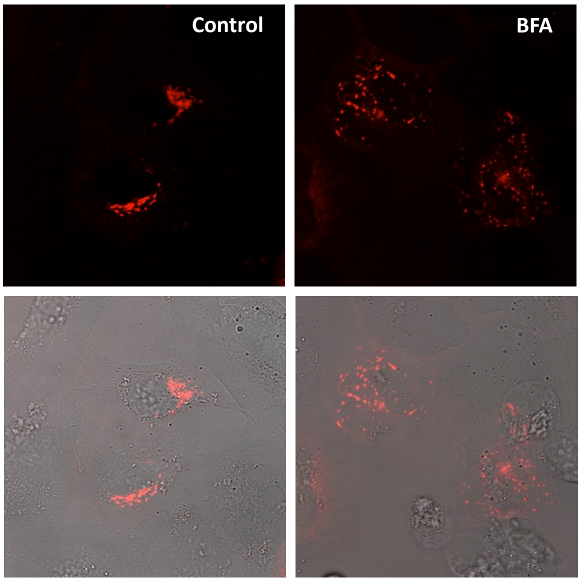


**Supplementary Figure 8. Imaging of Golgi apparatus in cells treated with BFA**. U2OS cells were transfected with mCherry-Giantin plasmid before (Control) and after (BFA) treatment with Brefeldin A (20 µM, 4 hours). Red: Golgi.

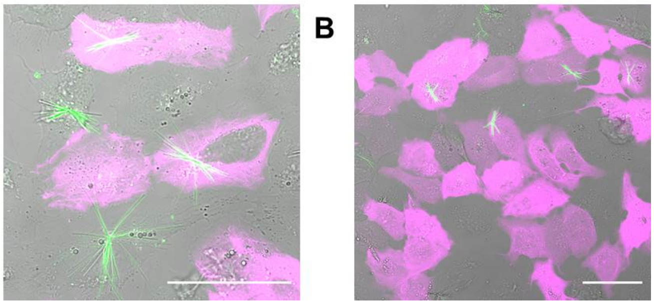


**Supplemenraty Figure 9. Imaging of transfected truncated kinesin in QPD-OTf treated U2OS**. A) Merge image of PTK2-GFP-Tubulin transfected with kin330 plasmid and treated with QPD-OTf (20 µM, 2.5 hours); green: crystals, magenta: kinesin. B) Merge image of PTK2-GFP-Tubulin transfected with kin560 plasmid and treated with QPD-OTf (20 µM, 2.5 hours); green: crystals, magenta: kinesin. Scale bar: 50 µm.

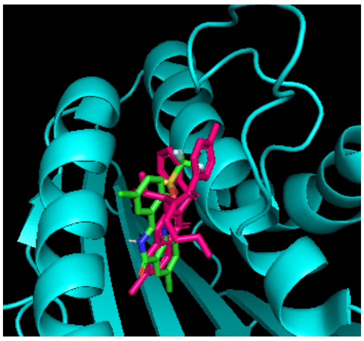

**Supplementary Figure 10**. **QPD-OTf docking into the Ispinesib binding site of Eg-5**. Superposition of Ispinesib (magenta) and QPD-OTf (green).

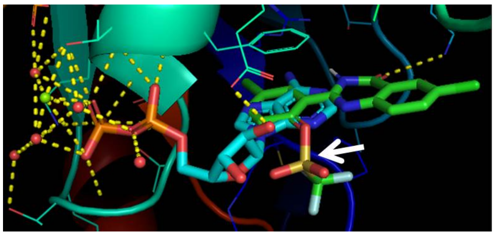

**Supplementary Figure 11. QPD-OTf docking into the ATP binding site of Eg-5**. Superposition of QPD-OTf (green) and ADP (cyan). White arrow indicates the triflate group of QPD-OTf.

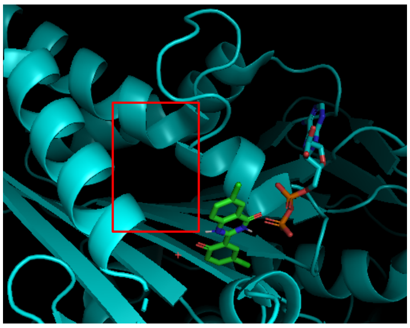

**Supplementary Figure 12. QPD-OTf docking into the Ispinesib binding site of kinesin-1 co-crystallized with ADP**. QPD-OTf (green), ADP (cyan). Red box indicates the “Ispinesib-like” binding site.

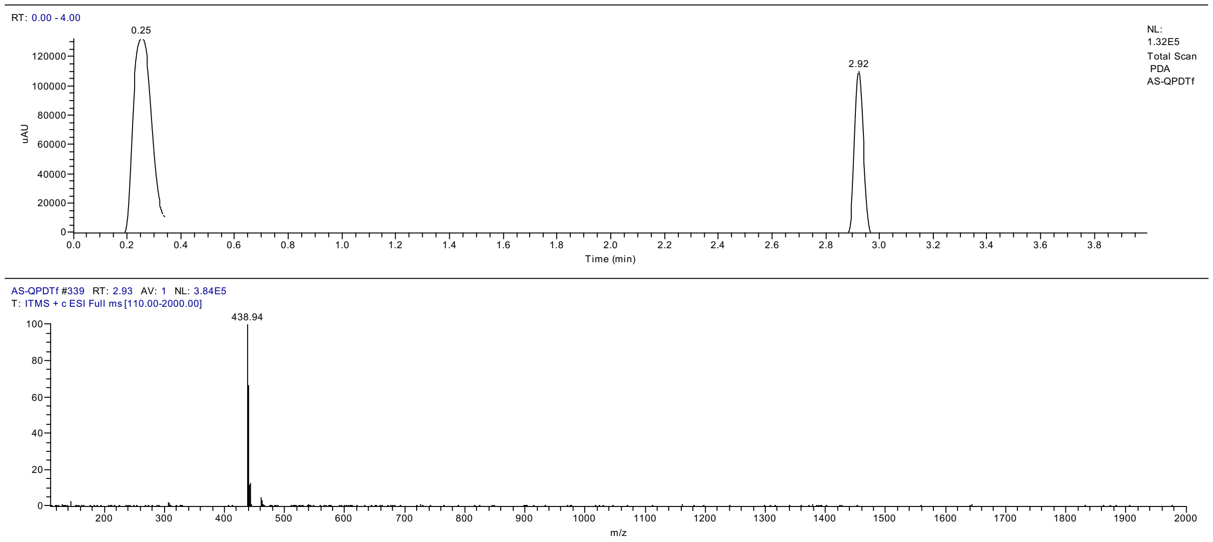


**Supplementary Figure 13**. LC-MS trace of QPD-OTf.

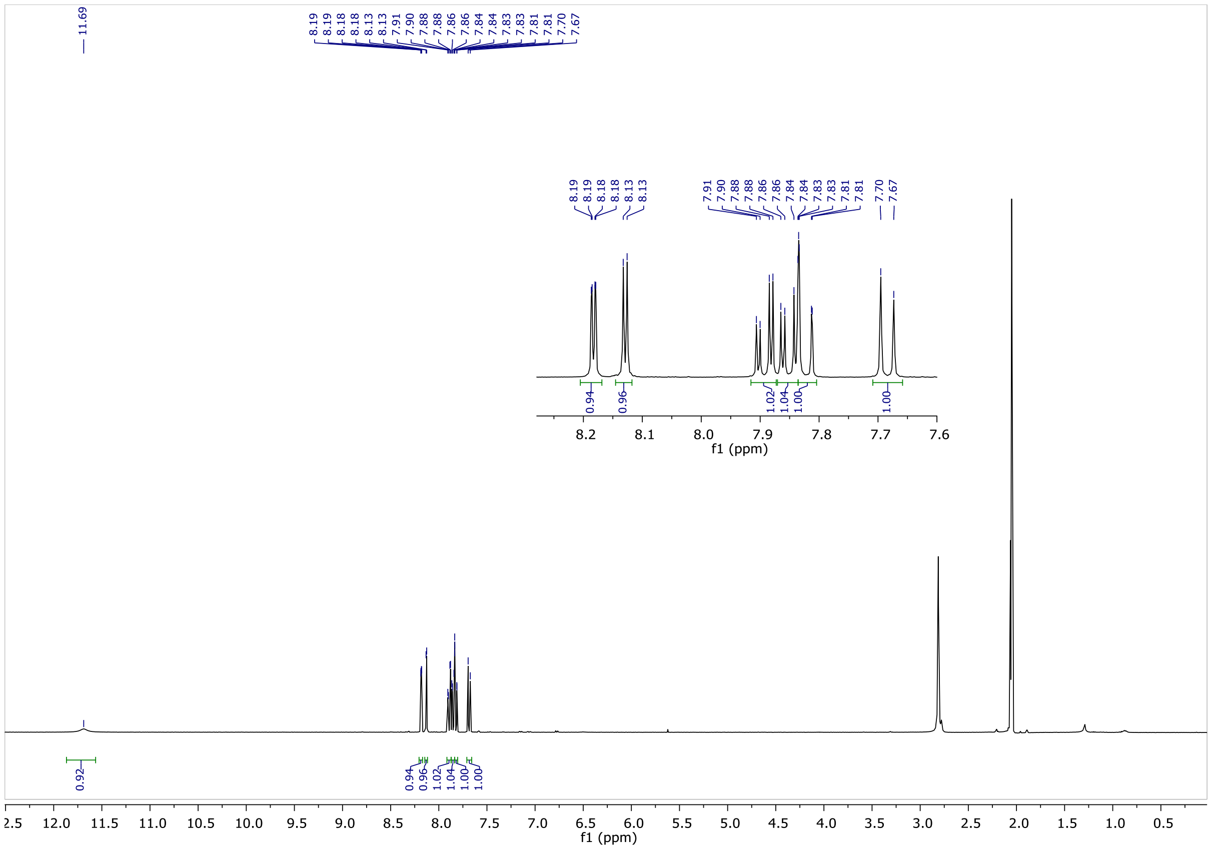


**Supplementary Figure 14**. $^{1}$H NMR of QPD-OTf

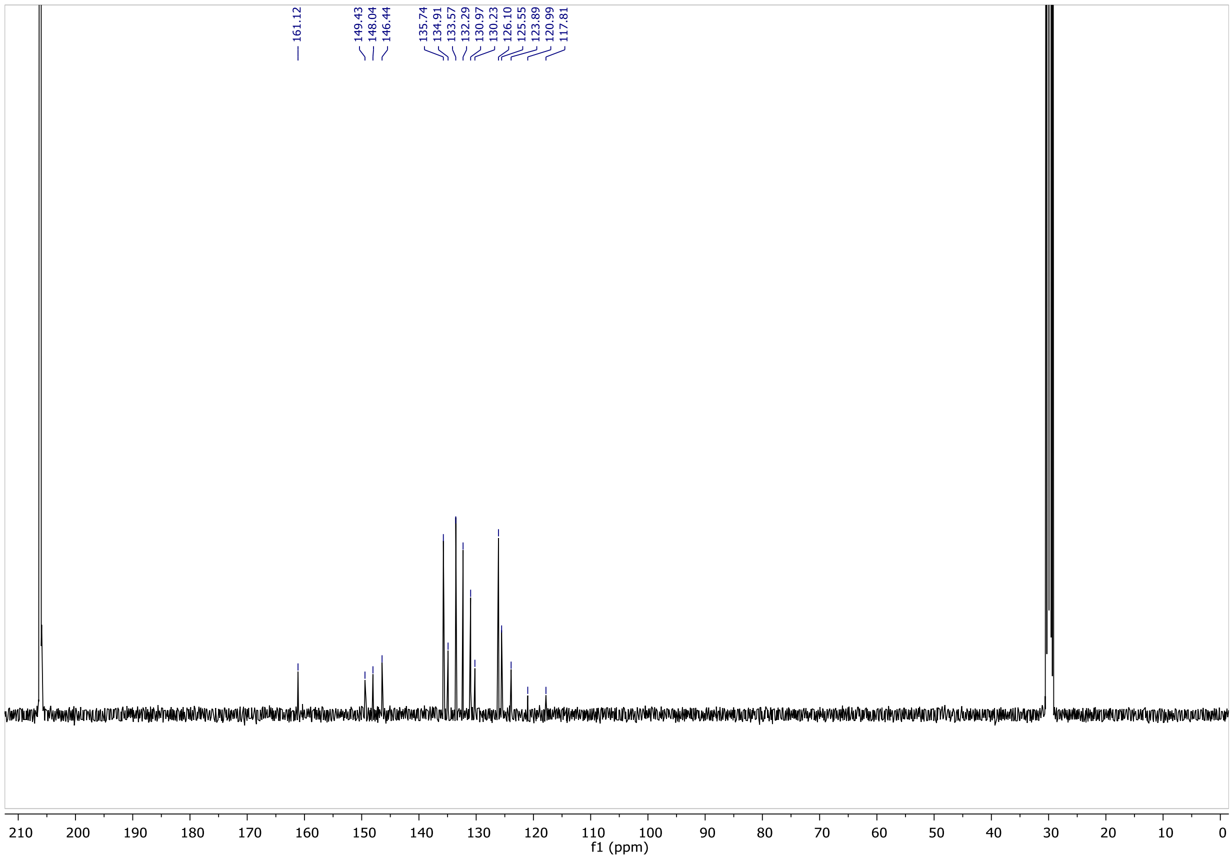


**Supplementary Figure 15**. $^{13}$C NMR of QPD-OTf

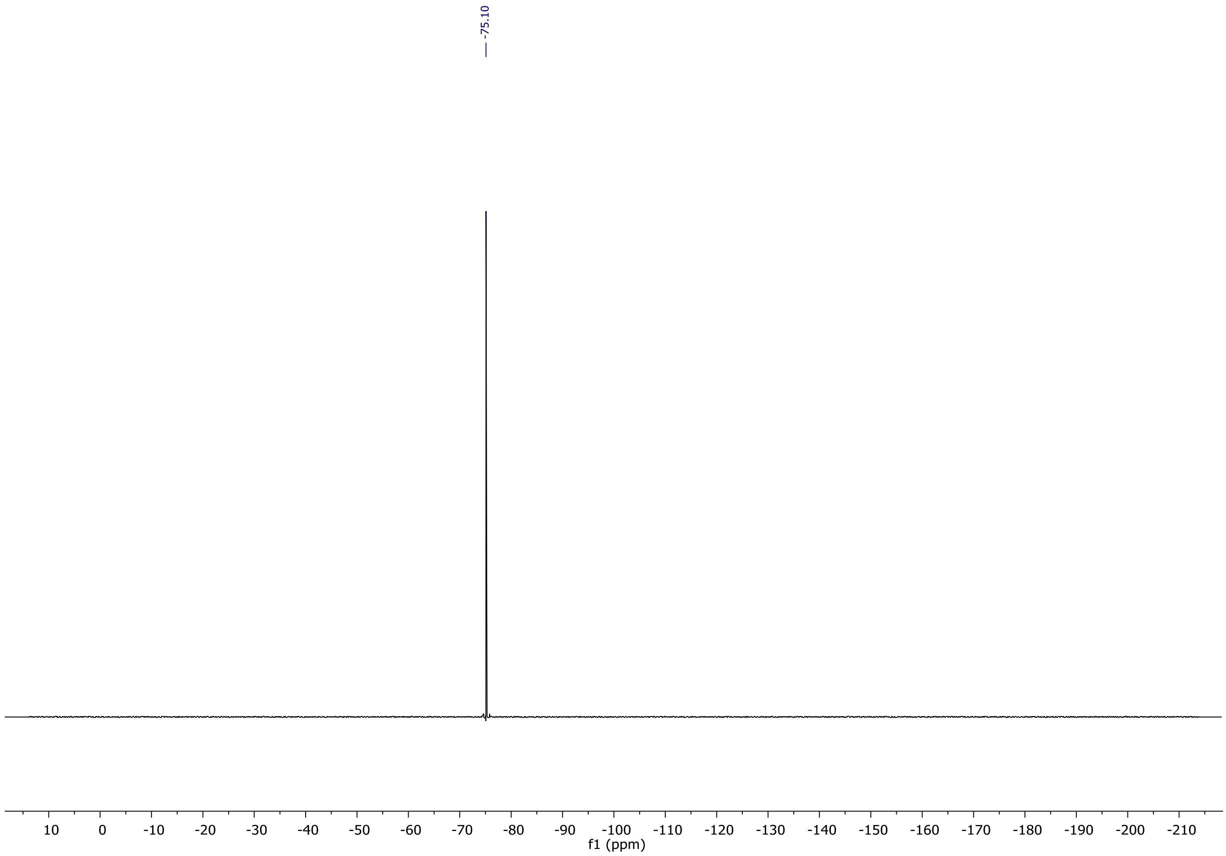


**Supplementary Figure 16**. $^{19}$F NMR of QPD-OTf.

### Supplementary Methods

#### Materials and methods

Chemical reactions were performed in anhydrous conditions under $N_2$. All reagents and solvents were purchased from commercial sources and were used without any further purification. Anhydrous solvents were obtained by passing them through commercially available alumina column (Innovative Technology, Inc., ® VA). Synthesized compounds were characterized using $^{1}H$, $^{13}C$, $^{19}F$ NMR ,recorded on AVANCE 3 HD for 400 MHz using Acetone-$d_6$, as solvent, with residual solvent peaks ($\delta$ = 2.05 ppm; $\delta$ = 29.84 ppm). LC-MS spectra were recorded by using a DIONEX Ultimate 3000 UHPLC coupled with a Thermo LCQ Fleet Mass Spectrometer System (electrospray ionization (ESI)) operated in positive mode (condition for elution gradient: 0 min, A:B = 100:0; 4 min, A:B = 10:90; solution A: 0.01% aqueous TFA solution; solution B, 0.01 % TFA in HPLC grade acetonitrile; flow rate: 0.750 mL/min.). Transmitted light imaging was performed using an EVOS XL Core Imaging System. Fluorescence imaging was carried out using a Leica SP8, a Zeiss LSM700 or a Zeiss LSM710 2P microscope. All images were analyzed with Image J. FIB-SEM rendering was performed with Imaris. pSF-mCherry-SNAP-Giantin plasmid was a kind gift of Howard Riezman's lab (University of Geneva, Switzerland). Docking calculations were performed with Autodock Vina.

#### Cell culture

U2OS, HeLa, HEK293T, MCF-7, RAW246.7 cell lines were obtained from the American Type Culture Collection (ATCC) and cultured according to their instructions. U2OS cells were grown in McCoy's 5A (modified) medium (Gibco) containing 10% FCS and 1% pen-strep at 37 °C under 5% $CO_2$ in a humidified incubator. Stable expressing GFP-Tubulin Ptk2 cells (kind gift from Franck Perez) were cultured in alpha-MEM (Gibco) containing 10% FCS and 1% pen-strep at 37 °C under 5% $CO_2$ in a humidified incubator. Cells were regularly tested for mycoplasma contamination by staining with Hoechst 33342.

#### QPD-OTf in Zymosan stimulated RAW246.7 cells

RAW246.7 were seeded into 3.5 cm glass bottom dishes with 10 mm microwell (Mattek); cells were incubated in DMEM medium at 37 °C under 5% $CO_2$ in a humidified incubator for 16 hours. Then media was removed, cells were washed twice with DPBS ($Ca^{2+}$, $Mg^{2+}$) and IgG-opsonized Zymosan A particles (20 µL) + QPD-OTf (20 µM) was added to cells in DMEM (-) (no additives). Cells were incubated at 37 °C under 5% $CO_2$ and imaged at different time points with a Zeiss LSM710 2P microscope.

**Sample Preparation for FIB-SEM**

HeLa cells were grown on MatTek™ glass coverslips for 2 days. Cells were then washed x 3 with Hank's Balanced Salt solution and QPD-OTf (20 μM) in DMEM without serum was added. Cells were then incubated for 4 hours at 37°C with 5% $CO_2$. Cells were then washed with Hank's Balanced Salt solution, fixed, and processed as previously described[1] with some differences. After dehydration the MatTek™ glass coverslip was removed from the plastic by using propylene oxide. The removed glass coverslip was then rinsed in 100% ethanol followed by immersion in mixtures of Durcupan ACM and ethanol with the following ratios: 25/75 for 1.5 hours, 50:50 for 1.5 hours; 75/25 overnight. The sample was then immersed in 100% Durcupan ACM for 4-5 hours with replacement of fresh Durcupan every hour. The glass coverslip was then removed and excess Durcupan was removed using filter paper. The coverslip was then placed in an oven at 60 degrees Celsius for 10 minutes, after which the sample was placed vertically in a 50 mL Falcon tube in folded filter papers and centrifuged for 15 minutes at 37°C and 750 RCF. The glass coverslip was then placed in an oven at 60 degrees Celsius under vacuum and left to polymerize over 2 days. The sample was then placed on a sample stub by an adhesive carbon dot, sputter coated with 50 nm gold, and painted with silver paint, followed by drying under vacuum.

**FIB-SEM**

Datasets were acquired using a Zeiss Crossbeam 540 (Carl Zeiss Microscopy GmbH, Jena, Germany). Platinum and Carbon was deposited over the region of interest and the run was setup and controlled by Atlas software (Fibics) SEM settings: 1.5 kV; 2.5 nA; Milling probe: 300 pA. The Slice thickness and the voxel size was set to 5 nm. The total volume acquired was: 16.36 x 9.87 x 7.31 μm (XYZ) and 23.5 x 9.60 x 7.47 μm (XYZ).

**FIB-SEM Data Analysis, Segmentation and Rendering**

The FIB-SEM datasets were aligned using Atlas5 software (Fibics). The data was then imported into Fiji software[2] and binned 3X, to 15 x 15 x 15 nm isotropic voxels. Segmentation of structures of interest was performed using the Pixel Classification module in the Ilastik software package (Ilastik.org) [3]. The probability maps were then imported into Imaris (Bitplane.com)[4] and surfaces were generated around fully segmented structures. Images and videos were rendered using Imaris. Crystal cross-sections based on surface renderings were measured in Imaris.

**Fixed cells imaging of QPD-OTf treated cells**

U2OS cells were grown in DMEM medium + 10% FBS to 50% confluency on 12 mm glass slides (seeded the day prior). Cells were treated for 24 hours with QPD-OTf (20 uM). After 24 hours, cells were fixed with MeOH fixation at -20 degrees Celsius for 5 minutes. Then the coverslips were washed for 30 minutes in PBS. Primary antibody staining was performed with 1:1000 dilution of DM1-alpha raised in mouse (T6199) and 1:1000 phalloidin raised in rabbit for 1 hour. Coverslips were washed in PBS for 30 minutes. Secondary antibody staining was performed with 1:400 dilution of anti-mouse ALEXA-488 and 1:400 dilution of 1:400 anti-rabbit ALEXA-568. Coverslips were washed in PBS for 30 minutes. Then coverslips were placed over DABCO mounting medium containing DAPI and imaged with a LSM700 microscope

**Tubulin purification from bovine brain and tubulin labelling**

Tubulin was purified from fresh bovine brain by two cycle of polymerisation and depolymerisation as previously described (Castoldi & Popov, 2003). A first polymerisation-depolymerisation cycle was performed in High-Molarity PIPES buffer [1 M PIPES-KOH at pH 6.9, 10 mM $MgCl_2$, 20 mM EGTA, 1.5 mM ATP and 0.5 mM GTP] supplemented with 1:1 glycerol and Depolymerisation buffer [50 mM MES-HCl at pH 6.6 and 1 mM $CaCl_2$] respectively. A second polymerisation-depolymerisation cycle was then performed: polymerisation in High-Molarity PIPES buffer and depolymerisation in 0.25XBRB80 complete after 15 min with 5XBRB80 to reach 1XBRB80 [80 mM PIPES at pH 6.8, 1 mM MgCl2 and 1 mM EGTA] respectively.

Labelled tubulin with ATTO-488, ATTO-565, or ATTO-647 (ATTO-TEC GmbH) and biotinylated tubulin were prepared as previously described (Hyman et al., 1991) with slight modification. Tubulin was polymerised in Glycerol PB solution [80 mM PIPES-KOH at pH 6.8, 5 mM $MgCl_2$, 1 mM EGTA, 1 mM GTP and 33 % (v/v) glycerol] for 30 min at 37°C and layered onto cushions of 0.1 M NaHEPES at pH 8.6, 1 mM $MgCl_2$, 1 mM EGTA and 60 % (v/v) glycerol followed by centrifugation. Pellet was resuspended in Resuspension buffer [0.1 M NaHEPES at pH 8.6, 1 mM $MgCl_2$, 1 mM EGTA, 40% (v/v) glycerol] and incubated 10 min at 37°C with 1/10 volume of 100 mM ATTO-488, -565, or -647 NHS-fluorochrome or incubated 20 min at 37°C with 2 mM Biotin reagent. Labelled tubulin was sedimented onto cushions of BRB80 supplemented with 60 % glycerol, resuspended in BRB80, and a second polymerisation-depolymerisation cycle was performed before use. Labelling ratio was 13 % for ATTO-565.

**rKin430-GFP expression and purification**

All *in vitro* experiments were performed with truncated, EGFP-labelled kinesin-1 construct, rKin430-EGFP (referred as kinesin-1 in the text). The rKin430-GFP plasmid (a kind gift of Stefan Diez's laboratory) was expressed and purified as previously described (Rogers et al., 2001). *E.coli* BL21(DE3)[pLysS] expressing rKin430-GFP were lysed in a Lysis buffer [50 mM Na-Phosphate buffer at pH 7.5, 300 mM KCl, 10 % Glycerol, 1 mM $MgCl_2$, 20 mM ß-Mercaptoethanol, 0.2 mM ATP, 30 mM imidazole and protease inhibitors cocktail tablets (Roche)]. The cleared lysate was loaded into a pre-equilibrated HisTrap column (GE Healthcare 1mL HisTrap column) using an ÄKTA Pure Protein Purification System (GE Healthcare). After predefined washes, protein was eluted with Elution buffer [50 mM Na-Phosphate buffer at pH 7.5, 300 mM KCl, 10 % Glycerol, 1 mM $MgCl_2$, 0.2 mM ATP, 300 mM imidazole and 10 % (w/v) sucrose]. A dialysis was performed overnight to exchange the Elution buffer 20 mM NaHEPES at pH 7.7, 150 mM KCl, 1 mM $MgCl_2$, 0.05 mM ATP, 1 mM DTT and 20 % (w/v) Sucrose. Protein concentration was measured by Bradford method and the concentration was adjusted to 1.2 µg/µL using Centrifugal filter Amicon 30K (Millipore). Protein was aliquoted and stored in liquid nitrogen.

**TIRF Imaging**

Microscopy imaging was realized with an Axio Observer Inverted TIRF microscope (Zeiss, 3i) and a Prime BSI (Photometrics). A 100X objective (Zeiss, Plan-Apochromat 100X/1.46 oil DIC (UV) VIS-IR) were used. SlideBook 6 X64 software (version 6.0.17) was used to record time-lapse imaging. For *in vitro*, microscope stage conditions were controlled with the Chamlide Live Cell Instrument incubator (37 °C).

**Flow chamber**

Slides and coverslips were cleaned by two successive incubations and sonication: sonicated for 40 min in 1 M NaOH, rinsed in bidistillated water, sonicated in ethanol (96 %) for 30 min and rinsed in bidistillated water. Slides and coverslips were dried with an air gun, placed into a Plasma cleaner (Electronic Diener, Plasma surface technology) for plasma treatment, followed by 2 days incubation with tri-ethoxy-silane-PEG (Creative PEGWorks) or a 1:5 mix of tri-ethoxy-silane-PEG-biotin and tri-ethoxy-silane-PEG at 1 mg/ml in 96 % ethanol and 0.02 % HCl, with gentle agitation at room temperature. Slides and coverslips were then washed in ethanol (96 %), and bidistillated water, dried with air gun and stored at 4 °C. Flow chamber was assembled by fixing with double tap a tri-ethoxy-silane-PEG treated slide with a 1:5 mix of tri-ethoxy-silane-PEG-biotin and tri-ethoxy-silane-PEG treated coverslip.

Microtubule seeds were prepared at 10 µM tubulin concentration (20 % ATTO-647-labelled tubulin and 80 % biotinylated tubulin) in BRB80 supplemented with 0.5 mM GMPCPP (Jena Bioscience) for 1 hour at 37°C. Seeds were incubated with 1 µM Paclitaxel (Sigma) for 45 min at 37 °C, centrifuged (50.000 rpm at 25°C for 15 min), resuspended in BRB80 supplemented with 1 µM Paclitaxel and 0.5 mM GMPCPP and stored in liquid nitrogen.

To observe precipitation of QPD in presence of stabilised microtubule seeds, we injected seeds in BRB80 supplemented with 20 µM QPD, 0.2 % BSA and anti-bleaching buffer (10 mM DTT, 0.3 mg/mL glucose, 0.1 mg/mL glucose oxidase, 0.02 mg/mL catalase, 0.125 % methyl cellulose (1500 cP, Sigma), 1 mM GTP, 2.7 mM $MgCl_2$ and 2.7 mM AMP-PMP), inside the flow chamber. No precipitation was observed after 2 hours under the microscope.

To study the precipitation of QPD in presence of dynamic microtubules we polymerised microtubules from seeds in BRB80 supplemented with 14 µm unlabelled tubulin, 20 µM QPD and 1 mM GTP at 37 °C in an Eppendorf-tube. For direct observation of precipitation, we polymerised microtubules from microtubule seeds in the flow chamber. The Flow chamber was prepared by injecting successively 50 µg/mL neutravidin (ThermoFisher), BRB80, GMPCPP microtubule seeds, and washed with BRB80 to remove unattached seeds. Microtubule polymerised from the seeds with 12 µM tubulin (20 % Atto-565 labelled) in BRB80 supplemented with an anti-bleaching buffer [10 mM DTT, 0.3 mg/mL glucose, 0.1 mg/mL glucose oxidase, 0.02 mg/mL catalase, 0.125 % methyl cellulose (1500 cP, Sigma), 1 mM GTP] and 0.2% BSA. The chamber was incubated for 15 min at 37 °C for polymerisation. The chamber sealed for imaging of microtubule dynamics and QPD precipitation at 37 °C of microtubule dynamics. Images were recorded every 3 min for 2 hours.

**Kinesore treatment in live cells**

U2OS cells (1.5x$10^5$) were seeded into 3.5 cm glass bottom dishes with 10 mm microwell (Mattek); cells were incubated in culture medium at 37 °C under 5% $CO_2$ in a humidified incubator for 24 hours. Then media was removed, cells were washed twice with DMEM (-) (no additives), Kinesore (100 µM) was added to cells in Ringer's buffer and cells were incubated at 37 °C under 0% $CO_2$ for 1.5 hours. Cells were imaged with a LEICA SP8.

**Molecular docking**

Docking calculations were performed with Autodock Vina.[5] Receptor (PDB structure: 3J8Y for kinesin-1, 4AP0 for Eg5) and ligand preparation were performed in AutodockTools. Results were displayed with PyMol.

**Synthesis of 4-chloro-2-formylphenyl trifluoromethanesulfonate**

5-chloro-2-hydroxybenzaldehyde (300 mg, 1.92 mmol, 1 eq) was dissolved in dry DCM (5 mL) and cooled to 0 °C and trimethylamine (535 µL, 3.84 mmol, 2 eq) was added. After 5 min triflic anhydride (320 µL, 1.92 mmol, 1 eq) was added dropwise and the mixture was stirred at 0 °C for 2 hours. Then the mixture was diluted with $H_2O$ (50 mL) and extracted with AcOEt (3x10 mL). The organic layers were combined, washed with brine, dried over $Na_2SO_4$ and concentrated under reduced pressure. The crude product was used in the next step without further purifications.
